# Learning Chirality-Aware Representations to Predict Drug Side Effect Frequencies

**DOI:** 10.64898/2026.05.14.725209

**Authors:** Aldo Galeano, Iago Dutra, Santiago Ferreyra, Alberto Paccanaro

**Affiliations:** Escola de Matemática Aplicada, Fundação Getulio Vargas, Praia de Botafogo, 22250-900, Rio de Janeiro, Brazil; Department of Computer Science, Centre for Systems and Synthetic Biology, Royal Holloway, University of London, Egham Hill, TW20 0EX, Egham, UK

**Keywords:** representation learning, drug side effects, graph neural networks

## Abstract

Ab initio prediction of side effect frequencies is important for assessing the risk–benefit profile of drugs and for identifying potential adverse effects early in development. A key challenge is chirality: many drugs exist as enantiomers, pairs of molecules with the same atoms and bond connectivity but different three-dimensional arrangements. Although chemically similar, enantiomers can interact differently with biological targets and therefore exhibit distinct efficacy and adverse-effect profiles. Here we introduce F2S (Features to Signatures), a method to predict the frequencies of drug side effects while explicitly accounting for chirality. Drug representations are learned directly from chemical structure using a directed-bond message-passing graph neural network that captures stereochemical configurations. Side effect representations are derived from curated textual descriptions encoded with a frozen PubMedBERT model. Side effect frequencies are predicted from the dot product between drug and side effect signatures together with biases for drugs and side effects. We evaluated F2S extensively across multiple settings, including cold-start and warm-start prediction, prospective evaluation, and scenarios controlling for chemical similarity between training and test drugs. Across these evaluations, F2S achieves performance comparable to state-of-the-art methods for general side-effect frequency prediction while producing fewer false positives and substantially improves the prediction of frequency differences between enantiomer pairs. Finally, F2S learns compact 10-dimensional signatures that support interpretability: drug signatures reflect therapeutic class and shared targets, side-effect signatures capture phenotype similarity, and the learned bias terms correlate with the popularity of drugs and side effects.

## Introduction

Adverse drug reactions are a major public health concern and a frequent cause of patient morbidity and mortality (Classen et al., 1997). They account for a substantial number of hospital admissions, impose a significant burden on healthcare systems, and in some cases lead to the withdrawal of drugs from the market (Onakpoya et al., 2016). Predicting adverse effects early in drug development is therefore an important goal in drug safety and pharmacology.

A meaningful assessment of the risk–benefit profile of a drug requires going beyond binary side effect prediction and estimating how frequently side effects occur. This problem was first addressed by a method developed in our lab (Galeano et al., 2020), which showed that side effect frequencies can be modelled through latent representations of drugs and side effects. We showed that it is possible to learn a signature for each drug and side effect such that the frequency of a side effect for a given drug can be inferred from the dot product between the corresponding drug and side effect signatures. Furthermore, we found that these signatures are linked to underlying biological processes in the cell. However, in that work, signatures for drugs and side effects were learned using a matrix decomposition approach that relied on known associations between drugs and side effects. As a consequence, making a prediction for a drug required at least a small number of observed side effects for that drug. This limitation is known in the recommender system literature as the “cold start” problem.

A seminal subsequent work, DSGAT (Xu et al., 2022), addressed this issue by learning drug signatures directly from chemical structures rather than from known side effects. In this approach, drug molecular graphs are encoded using graph neural networks to derive drug representations, which are then combined with learned side effect representations to predict side effect frequencies. This design enables the prediction for compounds with no previously observed side effects.

Until now, no method has addressed molecular chirality, a specific form of stereoisomerism that is an aspect of molecular structure highly relevant in drug development. A chiral molecule exists in two forms, known as enantiomers, which have the same chemical composition and bond connectivity but differ in their three-dimensional spatial arrangement (Neuman, 2013). These two forms are mirror images of each other and cannot be superimposed by any rotation, as illustrated in Figure 1. Although enantiomers are chemically similar, they can interact very differently with biological systems. As a result, one enantiomer of a drug may be therapeutically effective, while the other may be inactive or even cause adverse effects (Bernal and Lalancette, 2026). In DSGAT, drug representations are derived from chemical graphs that encode atomic composition and connectivity, but do not distinguish between enantiomers. Consequently, enantiomeric forms of a drug are mapped to identical representations, leading the model to predict exactly the same side effect profile and frequency for both forms.

**Figure 1.**
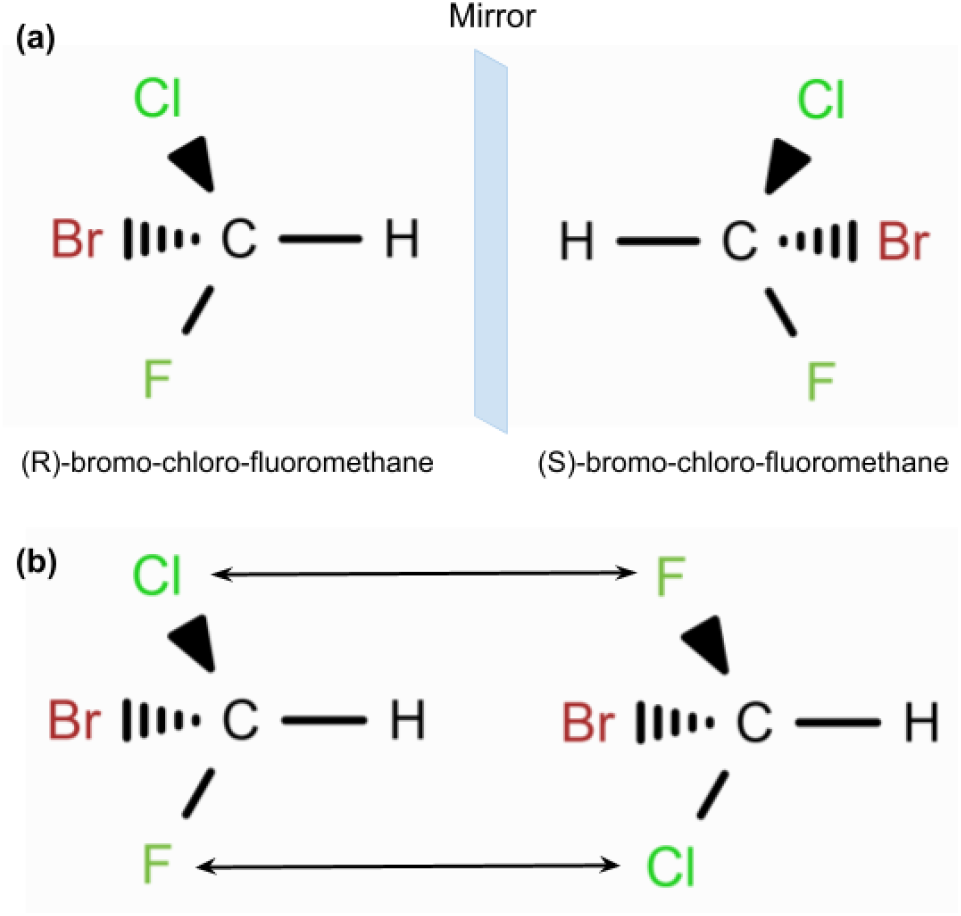
Optical isomerism. Bold wedges represent bonds coming above the image plane, and dashed wedges represent bonds going behind the image plane. **(a)** The R and S forms of bromo-chloro-fluoromethane are specular images of one another. **(b)** No rotation exists that can make them superimposed.

In this paper, we introduce a method that, to our knowledge, is the first for predicting the frequencies of drug side effects that directly addresses this limitation. Our approach learns drug signatures from chemical structure while explicitly incorporating chirality information, thus enabling the model to distinguish between enantiomeric forms of the same compound. At the same time, side effect signatures are learned from textual descriptions, allowing drug and side effect representations to be embedded in a shared latent space. As in earlier frequency-based models, predictions are obtained by the dot product between drug and side effect signatures. Our method achieves performance comparable to DSGAT on standard evaluation settings, while substantially improving accuracy in distinguishing enantiomeric pairs. Importantly, it does so using substantially lower-dimensional representations, and our experiments show that this compact representation supports a degree of interpretability in the learned features.

## Methods

### Dataset

Adapted the dataset curated by Galeano (Galeano et al., 2020), which contains 759 drugs and 994 side effects organized as a matrix (*R*) whose values were taken from SIDER (Kuhn et al., 2016). For each possible drug-side effect pair, a score of 0 indicates no known association between a drug and a side effect. Scores from 1 to 5 (1 = very rare, 2 = rare, 3 = infrequent, 4 = frequent, 5 = very frequent) represent the frequency class of a side effect, according to the standardized frequency categories established by the Council for International Organizations of Medical Sciences (CIOMS).

Ten protein-based drugs were discarded, as their length is computationally intractable for our method. Consequently, the adapted dataset comprises *D* = 749 drugs and *S* = 994 side effects, leading to a frequency matrix *R* containing 37,040 frequency scores. For each drug, isomeric SMILES were retrieved from PubChem (Kim et al., 2018). These isomeric SMILES are essential because they encode the chiral configuration of the bonds in the molecule. The complete dataset can be found in GitHub, along with the rest of the code.

### Model Architecture

In our earlier work, we showed it is possible to learn a non-negative embedding space such that the frequency scores can be estimated by the dot-product between drug and side effect representations (Galeano et al., 2020). Our aim is to improve on this work by obtaining the representation for drugs directly from the chemical structure, while relying on textual descriptions to obtain side effect representations. This would allow for the prediction for completely new drugs and side effects. Moreover, we extend the dot-product structure by incorporating explicit bias terms for drugs and side effects, as well as a global bias term. Such bias terms have been shown in the recommender systems literature to incorporate the popularity of an item (Koren et al., 2021). In our setting, we expect the model to infer these biases directly from the initial representations of drugs and side effects, so that they may capture certain tendencies, such as whether a drug is generally more prone to causing side effects or whether a side effect is more prevalent across drugs.

Figure 2 summarizes our model, named F2S (Features to Signatures). RDKit https://www.rdkit.org was used to build molecular graphs and extract initial chemical features from the drugs’ isomeric SMILES. The resulting graphs and initial chemical features are then processed by a message-passing graph neural network (GNN). In the last layer of the GNN, a global aggregation is performed to produce an embedding for the drug. This is scaled and transformed by a small Multi-Layer Perceptron (MLP) to create the final drug representation 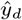 and scalar bias term *b*_*d*_. Simultaneously, the textual descriptions of side effects are processed using a frozen instance of PubMedBERT (Gu et al., 2021). Liu et al. collected these descriptions from Wikipedia and PubMed and curated them to avoid data leakage, for example by masking sentences that mention drugs (Liu et al., 2021). The 768-dimensional descriptor vector from PubMedBERT is reduced via Principal Component Analysis (PCA) to 64 dimensions and passed to a MLP block that produces the final side effect representation 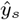 and bias *b*_*s*_. Predictions are made by summing the dot-product of drug and side effect representations with their respective bias terms and a global average *µ*. In our experiments, *µ* was always set to the average score of the training examples, and the size of the final representations was set to 10 dimensions.

**Figure 2.**
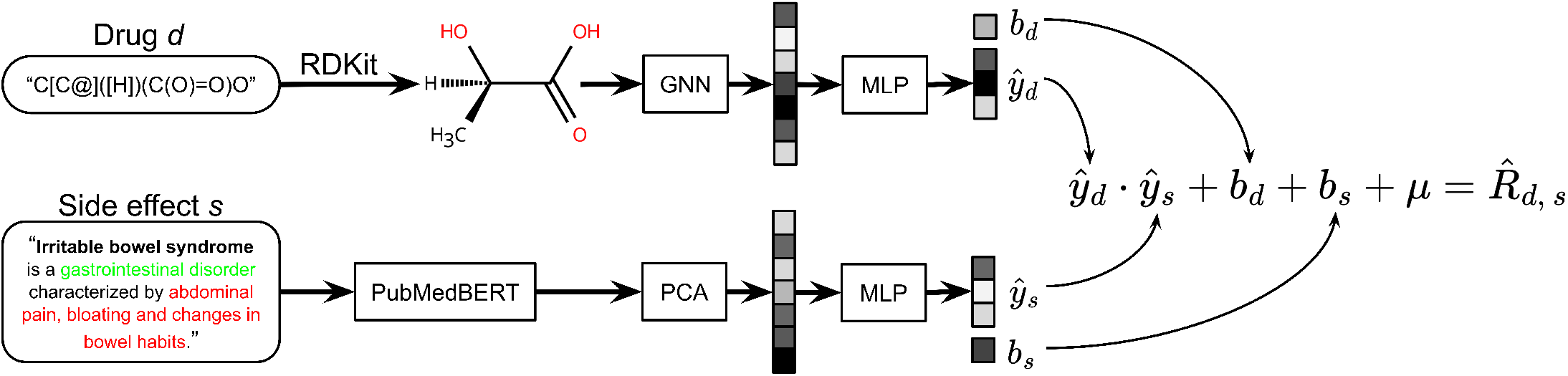
F2S architecture. Drugs stored as isomeric SMILES strings are transformed into molecular graphs and processed by a GNN module. After the global pooling step, a MLP head produces the final drug representation 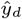 and drug bias *b*_*d*_. Side effects textual descriptions are first processed by PubMedBERT. The output is compressed via PCA, and then transformed by a MLP, resulting in a side effect representation 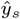 and a side effect bias *b*_*s*_. The final prediction consists on the sum of the dot-product between representations and drug and side effect specific biases with the global bias *µ*.

We now describe the GNN component of the model in more detail. The GNN is constructed to capture fine-grained structural patterns by performing directed message passing on the line graph derived from the original molecular graph. Given a molecule, we first construct an undirected graph *𝒢* = (*𝒱, ℰ*), in which atoms are represented as nodes and covalent bonds as edges. Each atom *i ∈* 𝒱 is associated with a feature vector 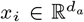, while each undirected bond (*i, j*) ∈ *E* is described by a bond feature vector 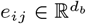. We then transform this undirected structure into a directed bond graph. For every bond (*i, j*), we introduce two directed edges, *i* → *j* and *j* → *i*. This construction allows information to propagate asymmetrically between neighboring atoms, which is crucial for distinguishing stereochemical configurations.

The initial representation of each directed bond is obtained by combining the features of the source atom with the corresponding bond features. Formally, we define

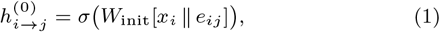

where ∥ denotes vector concatenation, *W*_init_ is a learnable parameter matrix, and *σ*(*·*) is a nonlinear activation function, such as ReLU. In this way, each directed edge embedding encodes both atomic identity and bond-specific information from the outset.

Message passing is then performed over the directed bond graph for *T* iterations. A directed bond *i* → *j* receives information from other directed bonds that terminate at its source atom *i*. We therefore define its neighborhood as

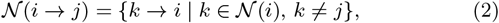

where *𝒩* (*i*) denotes the set of atoms bonded to *i*. Excluding *j* prevents immediate backtracking along the reverse edge. At iteration *t*, incoming messages are aggregated as

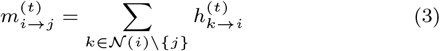

This aggregation step allows each directed bond to incorporate contextual information from its local chemical environment. The representation is then updated by combining the initial embedding with the aggregated message:

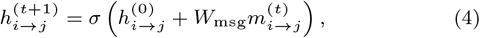

where *W*_msg_ is a learnable weight matrix shared across iterations. By repeatedly applying this update, the model progressively expands the receptive field of each directed bond representation, allowing it to encode higher-order structural dependencies.

After *T* message passing steps, we produce final node representations by fusing initial features with the learned chemical context. Specifically, we define

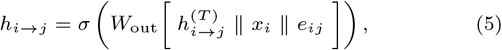

where *W*_out_ is a learnable parameter matrix.

A graph-level molecular representation is then obtained by mean pooling over all embeddings:

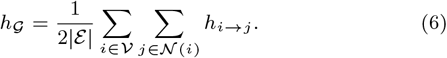

Finally, the aggregated representation *h*_*G*_ is passed through a MLP to produce the final molecular signature 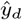. By propagating information along directed bonds and conditioning each update on the initial atom and bond features, this mechanism captures asymmetric local structural patterns. Consequently, the model becomes sensitive to subtle geometric differences induced by isomerism, improving its ability to distinguish stereochemical variations that may lead to distinct biological effects.

### Training

Our model is trained by minimizing the loss function:

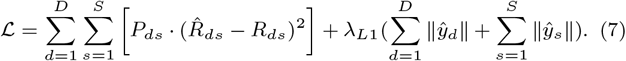

Similarly to our earlier work (Galeano et al., 2020), the loss function consists on the squared weighted Frobenius norm of the difference between predicted 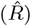 and real (*R*) frequency scores (Equation 7). The pondering matrix *P* is such that *P*_*ds*_ = *α* if *R*_*ds*_ = 0 and *P*_*ds*_ = 1 otherwise. The zeros are downweighted in order to handle the sparcity of *R* and to diminish the impact of unknown associations. Throughout our analyses, *α* was set to 0.03 in accordance with Galeano (Galeano et al., 2020). Additionally, to enforce sparcity in the learned representations, we penalize the L1-norm of drug and side effect embeddings, 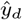 and 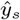, with a coefficient *λ*_*L*1_ that controls its importance. Furthermore, we apply weight decay as regularization on the norm of the model’s parameters.

The parameters were optimized using PyTorch’s implementation of Adam. The values for learning rate, weight decay and L1-regularization were decided by grid-search using cross-validation and were set to 1 *·* 10^*−*4^, 1 *·* 10^*−*5^ and 1 *·* 10^*−*4^, respectively.

## Results

The goal of F2S is to predict side effect frequencies for compounds for which none are known, and to predict different side effect profiles for a drug and its enantiomer when appropriate. We compared the performance of F2S against DSGAT (Xu et al., 2022), which, to our knowledge, is the only approach that relies exclusively on chemical features to represent drugs, enabling the analysis of novel compounds. In our experiments, we first evaluated the models’ predictions for novel compounds, and more specifically, for compounds that are enantiomers of drugs with known side effects. We further evaluated F2S by predicting the side effect frequencies for drugs with already known side effects, comparing its performance with DSGAT and Galeano’s approach (Galeano et al., 2020). In addition, we performed a prospective evaluation, where we assessed whether F2S could recover side effects that were discovered only after a drug entered the market.

This section also includes the results of an analysis of the signatures learned by F2S. The aim of this analysis was to determine whether the learned drug and side effect signatures capture intrinsic biological properties. Finally, we examined whether the learned drug and side effect bias terms are related to their respective popularities.

### A. Predicting frequencies of drug side effects

Our problem is a multi-class ordered classification task with five classes describing the frequency of side effects and one class representing the absence of side effects (denoted as 0 in the frequency matrix). As done previously by DSGAT and Galeano’s approach, we assigned classes to F2S predicted scores using a thresholding operation. To do this, we divided the predicted scores for each drug–side effect pair used in training into six sets (one set per frequency class, including the zero class). For each set, we estimated a probability density function of the scores using Gaussian kernel density estimation. Class thresholds were then determined via maximum likelihood, assigning each predicted score to the frequency class whose estimated density was highest.

The performance of a model in predicting each frequency class *I* can be evaluated using recall and precision. The recall for class *i* is defined as the proportion of side effects with frequency *i* in the testing set that are predicted by the model as belonging to class *i*. A high recall indicates that most instances belonging to class *i* are successfully identified. The precision for class *i* is the proportion of side effects predicted by the model as belonging to class *i* that indeed belong to that class in the testing set. A high precision indicates that most predictions assigned to class *i* are correct. These two metrics are complementary and can be summarized by the F1 score, which is the harmonic mean of precision and recall.

In our experiments, we computed the F1 score for each class and for each drug, and reported the mean F1 score per frequency class across all drugs. Additionally, we computed the mean across the F1 scores of all frequency classes, denoted as Macro-F1, to summarize the overall performance. This metric reflects the practical applicability of the models, as it jointly accounts for the proportion of correctly predicted side effect frequencies and the proportion of false positives among predicted frequencies (Sokolova et al., 2009).

#### Predicting side effects frequencies for enantiomers

We designed F2S to learn chirality-aware drug signatures, addressing the fact that two drugs with very similar molecular structures may cause different effects on the human body. In our dataset, we found eight pairs of drugs that are enantiomers of one another. Table 1 shows these drug pairs in the first column, while the second column reports the number of side effects for which they exhibit different frequencies in our dataset. Notably, approximately 50% of these differences correspond to cases in which one drug manifests a side effect that its enantiomer does not.

**Table 1.**
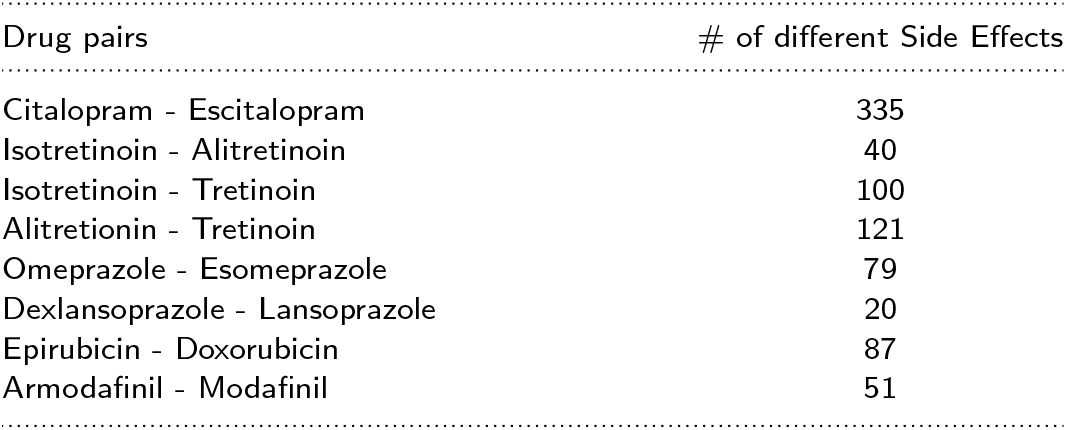
Side effect differences between enantiomers.

| Drug pairs | # of different Side Effects |
| --- | --- |
| Citalopram - Escitalopram | 335 |
| Isotretinoin - Alitretinoin | 40 |
| Isotretinoin - Tretinoin | 100 |
| Alitretinoin - Tretinoin | 121 |
| Omeprazole - Esomeprazole | 79 |
| Dexlansoprazole - Lansoprazole | 20 |
| Epirubicin - Doxorubicin | 87 |
| Armodafinil - Modafinil | 51 |

To investigate whether the models can capture these stereospecific differences, we evaluated F2S and DSGAT on their ability to predict the side-effect profile of a held-out enantiomer. We employed a “leave-one-enantiomer-out” strategy: Drug A (the training instance) was included in the training set, while its mirror-image counterpart, Drug B, was reserved for the test set. This setup specifically tests the models’ capacity to retrieve side effects unique to Drug B that are absent from the profile of Drug A.

The first column of Table 1 contains 13 drugs, therefore we trained each model 13 times. In each run, one drug was held out for testing, while all others were included in training. We then evaluated the models’ predictions for all side effects, excluding those shared between the held-out drug and its corresponding enantiomer. Figure 3 reports the mean F1 score for each frequency class across all drugs in Table 1, as well as the Macro-F1 score.

**Figure 3.**
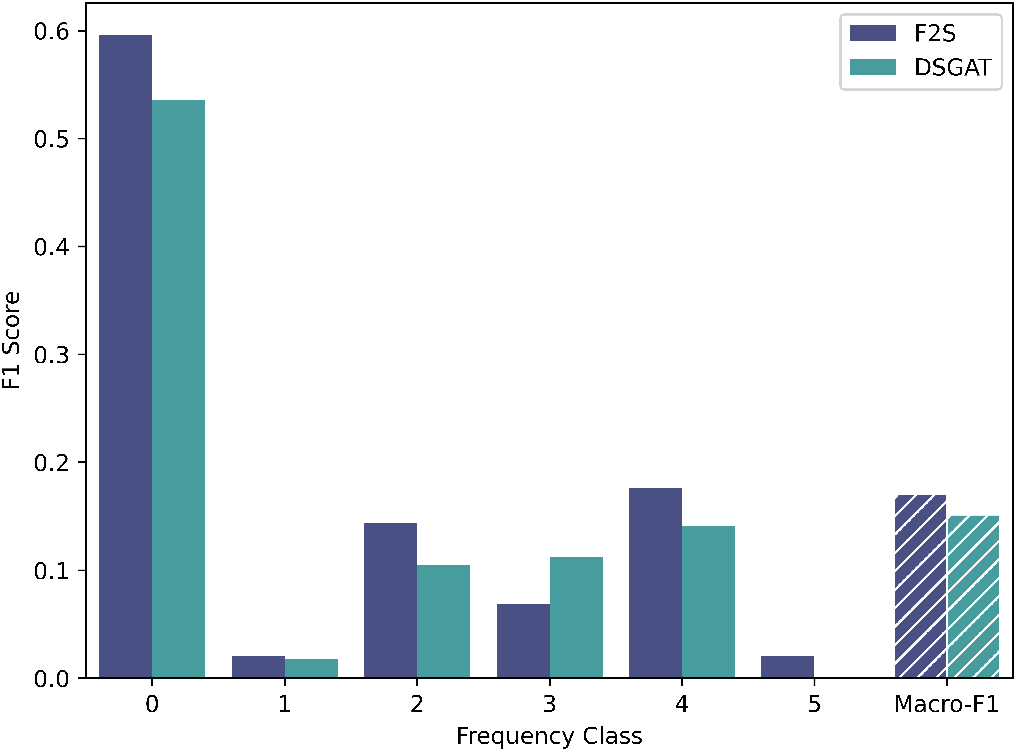
F1 Score for prediciting the side effect frequencies of enantiomers. Bar plot of the mean F1 score for each frequency class across the 13 enantiomeric drugs, along with the Macro-F1 score summarizing overall model performance.

F2S outperformed DSGAT in nearly all frequency classes. In particular, DSGAT failed to recover any side effects with frequency 5. This may reflect the fact that DSGAT assigns the same signature to drugs A and B, and the inclusion of Drug A in the training set effectively restricts the model to predict a similar side effect profile for Drug B. As a result, DSGAT is limited in its ability to assign high-frequency side effects to Drug B when those effects are absent for Drug A.

In contrast, F2S learns chirality-aware representations that allow enantiomers to occupy distinct positions in the learned signature space. This flexibility enables F2S to assign different side effect frequency profiles to drugs A and B.

#### Predicting side effects frequencies for novel compounds

We compared the overall performance of F2S and DSGAT at predicting side effect frequencies for new compounds, that is, compounds without any observed side effects in the training data. For this evaluation, we performed 35-fold cross-validation, where in each fold 21 or 22 drugs were left in the test set. These drugs were not observed during model training. Figure 4 shows a bar plot of the mean F1 score for each frequency class, along with the Macro-F1 score, averaged across all drugs. Overall, DSGAT and F2S exhibited comparable performance, with DSGAT achieving higher F1 scores for frequency classes 4 and 5. However, F2S produced significantly fewer false positives than DSGAT, suggesting that it more effectively distinguishes whether a drug can cause a given side effect. This resulted in a slightly higher Macro-F1 score across all drugs.

**Figure 4.**
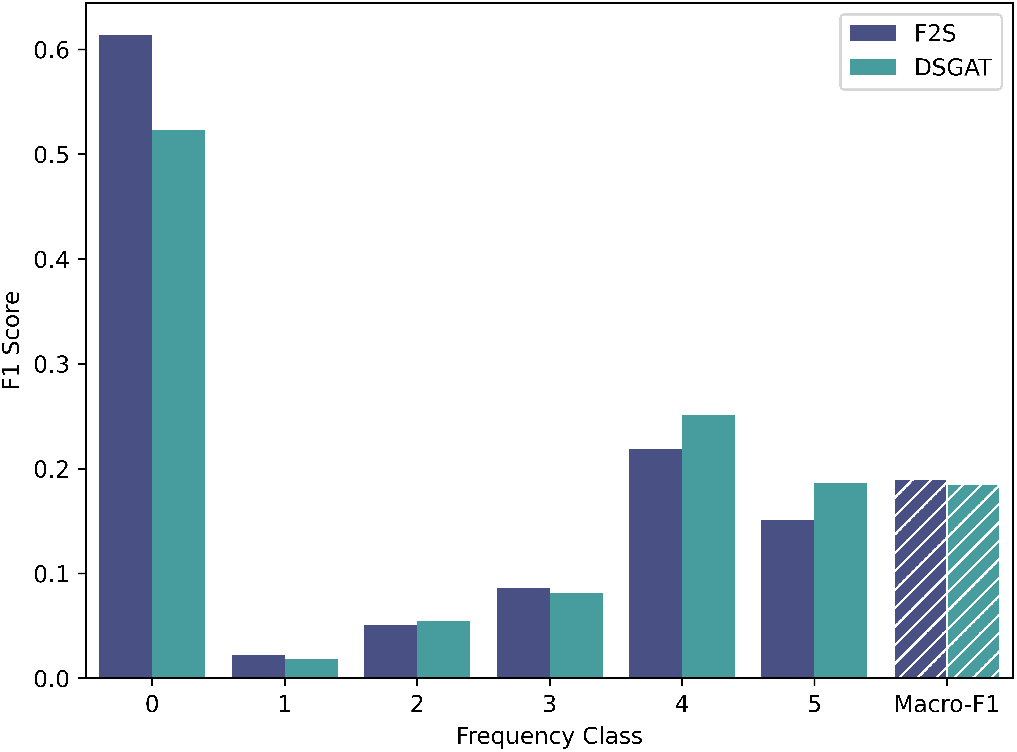
F1 Score for novel compounds. Bar plot of the mean F1 score for each frequency class across all drugs, along with the Macro-F1 score summarizing overall model performance.

Moreover, we assessed whether model performance is affected by high chemical similarity between drugs in the testing set and those in the training set, that is, whether the models were simply predicting similar side effect frequencies for test drugs that were chemically similar to drugs in the training set. To this end, we computed pairwise chemical similarity using the Tanimoto coefficient over Morgan fingerprints. We identified 215 drugs in the testing set whose maximum similarity to any drug in their respective training set was below 0.3 and used them as a low-similarity testing set.

Under this more challenging scenario, the mean Macro-F1 score of F2S decreases from 0.1905 to 0.1775. Despite this reduction, F2S retained substantial predictive capability even for chemically dissimilar drugs. Similarly, DSGAT’s performance decreased from 0.1856 to 0.1694. For frequency class 0 specifically, DSGAT’s mean F1 score decreases from 0.52 to 0.45, whereas F2S shows a smaller reduction, from 0.61 to 0.57. This comparatively smaller decline indicates that F2S is less sensitive to the removal of chemically similar training examples. Overall, these results suggest that F2S generalizes more robustly than DSGAT when predicting side effect frequencies for structurally dissimilar compounds.

#### Predicting side effects for known drugs

It is valuable to assess a model’s performance at predicting missing side effect frequencies for drugs with a set of already known side effect frequencies. For this evaluation, we randomly set to zero 10% of the non-zero entries of matrix *R*. The models were trained on this modified matrix, and the F1 score was computed for all frequency classes across all drugs on the held out set. The resulting Macro-F1 scores for Galeano’s method, DSGAT, and F2S are 0.1558, 0.1401 and 0.1368 respectively.

For frequency class 0, the mean F1 scores are 0.73, 0.63 and 0.64, respectively. Overall, F2S performs comparably to DSGAT in this setting, while Galeano’s method slightly outperforms both DSGAT and F2S. This may be due to its explicit use of the known side effect frequencies in training.

#### Predicting post-marketing side effects

Many side effects are identified only after a drug has entered the market. The ability of F2S to recover missing side effects motivated us to investigate whether it could also be used to predict previously unknown side effects. This setting can be viewed as a prospective evaluation: we trained F2S using side effect frequencies observed during clinical trials and subsequently assessed its ability to predict side effects that are identified after market approval.

We used two independent post-marketing test sets for this analysis from the SIDER and OFFSIDES datasets (Kuhn et al., 2016; Tatonetti et al., 2012). In our evaluation, we divided the predicted scores into three groups: (1) scores corresponding to no side effects in the test set; (2) scores corresponding to post-marketing side effects reported in SIDER; and (3) scores corresponding to post-marketing side effects reported in OFFSIDES. Using the Wilcoxon rank-sum test, we found that the scores associated with SIDER and OFFSIDES side effects were significantly higher than those associated with no side effects. Furthermore, since the associations reported in SIDER are generally considered more reliable than those in OFFSIDES (Galeano et al., 2020), we computed the AUROC for predicting SIDER side effects and obtained a value of 0.73. This result suggests that many post-marketing side effects in these datasets could have been detected by F2S.

### B. Analysis of F2S-learned signatures

F2S maps the chemical features of the drug and the textual information of the side effect encoded by PubMedBERT into a shared low-dimensional signature space for drugs and side effects. The strong predictive performance of F2S on new compounds suggests that these learned signatures may capture latent drug and side effect properties. To assess this hypothesis, we analyzed the structure of the learned signature space and verified the consistency of our findings in 30 runs of F2S.

#### Drug signatures are informative of main drug activity

We divided the drugs in our dataset into different groups according to their clinical activity. For this classification, we used the main Anatomical, Therapeutic, and Chemical (ATC) class levels (http://www.whocc.no/atc/structure_and_principles/). The ATC system is a hierarchical classification maintained by the World Health Organization, where terms at lower levels correspond to more specific descriptors of clinical activity. Each drug in matrix *R* is associated with its respective ATC category at three different levels. The drugs in this dataset cover all 14 groups at the general Anatomical level, 70 at the intermediate Therapeutic level, and 147 at the more specific Chemical level.

We analyzed whether the cosine similarity between drug signatures is informative of drugs belonging to the same ATC class. To this end, we tested whether similarities between drugs within the same ATC class are significantly higher than similarities between drugs from different classes across all 30 runs of F2S. Using the Wilcoxon rank-sum test, we found that this is indeed the case, indicating that the learned similarities can be used to group drugs according to their clinical activity.

We defined a binary classification task for each drug in which the signature similarity was used to predict which other drugs shared the same ATC class at that level. Figure 5a shows how the mean AUROC increases as we move to lower levels of the hierarchy, reflecting the fact that drug clinical activity becomes more similar at more specific levels.

**Figure 5.**
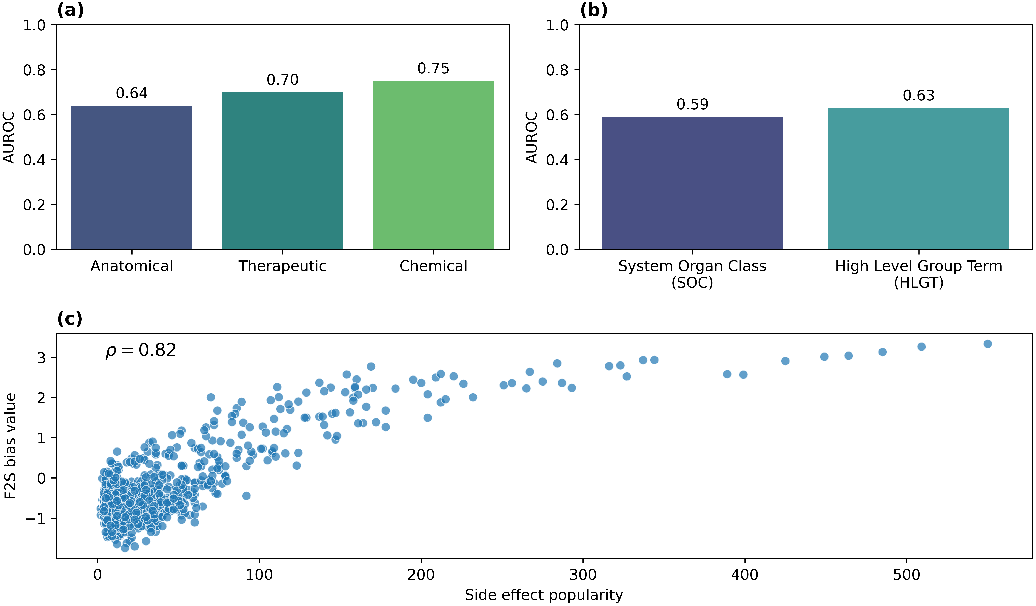
Interpretability of learned signatures. **(a)** AUROC when using drug signature cosine similarities to predict ATC categories. **(b)** AUROC when using side effect signature cosine similarities to predict MedDRA categories. **(c)** Correlation between learned side effect biases and side effects popularities.

We performed a similar analysis to assess whether the cosine similarity is informative of drugs targeting the same human protein. To this end, we obtained drug targets from Drug Bank (Knox et al., 2024). Across all 30 runs of F2S, the similarity between drugs targeting the same protein was statistically significantly higher than that between drugs targeting different proteins (Wilcoxon rank-sum test). The mean AUROC for this task was 0.75, further demonstrating that the signatures learned by F2S capture biologically meaningful information about drug effects.

#### Side effect signatures are informative of side effect phenotypes

We divided the side effects in our dataset according to their associated MedDRA (Medical Dictionary for Regulatory Activities) terms at two levels of its hierarchy (Brown et al., 1999). Our aim was to test whether side effects with high signature cosine similarity tend to share the same MedDRA terms. Across all 30 runs of F2S, the similarities between side effects sharing the same MedDRA term at both levels were statistically significantly higher than those between side effects belonging to different terms (Wilcoxon rank-sum test). Figure 5b shows how the mean AUROC increases as we move into more specific categorization.

#### The learned biases of drugs and side effects reflect their popularity

The “popularity of a drug” can be thought as the number of side effects associated with it. Analogously, the “popularity of a side effect” is the number of drugs that cause it. These quantities capture global prevalence effects in the dataset. To investigate whether the bias terms in F2S reflect such prevalence patterns, we computed the popularity measures for all drugs and side effects and compared them to the learned biases.

Specifically, we extracted the learned bias terms for all drugs and side effects across 30 independent runs of F2S and computed the correlation between the learned biases and the corresponding popularity measures. We observed a mean correlation of 0.7 for side effect biases and 0.4 for drug biases. These results indicate that F2S partially captures the prevalence of drugs and side effects through its bias terms. Figure 5c illustrates this trend for one run of the model, in which the Pearson correlation between side effect biases and popularities was 0.82.

## Discussion

In this paper, we present F2S, a method for predicting the frequencies of drug side effects from chemical structure. Unlike existing approaches such as DSGAT, which derive drug representations from molecular graphs but cannot distinguish enantiomers, F2S explicitly incorporates molecular chirality into the learning of drug signatures. As a result, it can learn distinct representations for enantiomeric forms of the same compound and predict different side-effect frequency profiles for each. Our experiments on enantiomeric pairs show that F2S recovers more side-effect frequencies across almost all frequency classes than DSGAT while producing fewer false positives.

An important aspect of F2S is that it preserves the simple linear relationship between drug and side-effect representations that we introduced previously (Galeano et al., 2020). In earlier work, we showed that side-effect frequencies can be predicted from the dot product between a drug signature and a side-effect signature, revealing a linear relationship between these representations and the observed frequencies. Similarly to DSGAT, F2S builds on this idea by learning drug signatures directly from chemical structure while retaining this linear prediction mechanism. In this sense, both approaches extend our original framework by learning embeddings from molecular features, rather than from known drug-side effect associations. Importantly, F2S further improves this framework by explicitly incorporating stereochemical information and distinguishing enantiomeric forms of the same compound.

Another aspect of F2S is that it is able to achieve competitive predictive performance even using very compact representations for drugs and side effects. In our experiments, the learned signatures have only 10 dimensions, while the results reported for DSGAT used representations of 200 dimensions. We were able to show that the learned representations nevertheless capture meaningful biological structure. Similarities between drug signatures correlate with ATC therapeutic categories and shared drug targets, while similarities between side-effect signatures reflect their organization in the MedDRA hierarchy. In addition, the learned bias terms correlate with the popularity of drugs and side effects, suggesting that the model captures global tendencies present in the data. The small number of dimensions may also facilitate the interpretation of individual latent factors in future work.

Predicting side-effect frequencies is naturally a multi-class classification problem, where the goal is to correctly recover the frequency category for each drug–side effect pair. For this reason, we evaluate model performance using the F1 score. In sparse settings such as ours, a model may achieve high recall for a given class by predicting many false positives, and the F1 score accounts for this trade-off between recall and precision. Other metrics such as root mean square error, mean average precision, and AUROC have also been used in this context and can provide useful information. However, when evaluated at the class level, these metrics do not penalize false positives in the same way. The F1 score therefore provides a more appropriate measure of performance in this multi-class, sparse prediction setting.

Future work will focus on further analysing the learned representations in order to better understand how specific molecular features influence predicted side-effect frequencies.

## Data availability

The data and code underlying this article are available in https://github.com/paccanarolab/Features2Signatures.

